# Probing the Ca_V_1.2 interactome in heart failure identifies a positive modulator of inotropy and lusitropy

**DOI:** 10.1101/2025.05.08.652977

**Authors:** Aaron Rodriques, Guoxia Liu, Alex Katchman, Sergey Zakharov, Elaine Wan, Marian Kalocsay, Robyn Eisert, Gary A. Bradshaw, Bi-Xing Chen, Lin Yang, Steven Reiken, Xianghai Lai, Ruiping Ji, Najla Saadallah, Veli Topkara, Stavros Fanourakis, Qi Yuan, Jared Kushner

**Author notes:** Corresponding author: Jared Kushner (Mailing Adress 630 W 168^th^ St., Presbyterian Hospital Building 8-405, New York, New York, 10032.

## Abstract

**Background:** Maladaptive changes in the function, expression and localization of proteins involved in calcium-handling worsen the impaired contractility of systolic heart failure (HF). Standard proteomics techniques require cell lysis and so are unable to characterize changes specific to the critical sub-cellular domain bounded by the T-tubule and the sarcoplasmic reticulum, known as the cardiac dyad. Traditional approaches are also less likely to capture low-affinity protein-protein interactions on lipid membranes. To improve our understanding of heart failure pathophysiology, we applied proximity proteomics to the cardiac dyad of mice with ischemic cardiomyopathy.

**Methods:** Using two lines of transgenic mice expressing fusion proteins of the engineered ascorbate peroxidase APEX2 with subunits of the cardiac voltage gated calcium channel, Ca_V_1.2, we labeled the dyad proteome of live, intact myocytes from healthy mice (N=6) and mice with coronary artery ligation HF (N=5) by peroxidase-catalyzed biotinylation. Quantitative mass spectrometry with isobaric tandem mass tags (TMT) was used to assess alterations in the local dyad proteome in myocytes from mice with chronic, remodeled HF. We subsequently generated a mouse with inducible cardiac overexpression of Galectin-1 to examine the effects of this protein on cardiac function and more specifically on the calcium-handling properties of mature myocytes, using cellular electrophysiology, calcium imaging, and echocardiography.

**Results:** From mice with HF, we found significant enrichment of 43 proteins defined by their abundance and proximity to transgenic Ca_V_1.2 α_1C_ channels, and a significant reduction in 22 proteins, out a of a total of 2326 proteins quantified. We also significantly enriched 286 proteins, and saw a reduction in 13 proteins, defined by proximity and abundance to Ca_V_1.2 β_2B_-subunits, out of a total 2236 proteins identified. Pathway analysis revealed HF is associated with increased abundance of components of the 26s proteasome and microtubules in the dyad, as well as the dimerizing, carbohydrate binding protein Galectin-1. Cardiac specific overexpression of Galectin-1 in healthy mice increases activation of Ca_V_1.2 and the Ryanodine receptor and accelerates myocyte relaxation through phosphorylation of Phospholamban.

**Conclusions:** Using proximity proteomics to examine the effects of HF *in vivo*, we find increased localization of Galectin-1 to the cardiac dyad, and that overexpression of Galectin-1 accelerates calcium kinetics in the heart.

## INTRODUCTION

Despite remarkable gains in recent years, heart failure (HF) remains the leading cause of morbidity and mortality in the US and the world^1^. Heart failure pathophysiology is characterized by multiple changes to the structure and function of cardiomyocytes. For instance, there is disorganization and rarefaction of the transverse tubule (T-tubule) network in systolic HF^2–4^. The cell membrane of T-tubules closely apposes the membrane of the sarcoplasmic reticulum (SR), where the calcium (Ca^2+^) that drives cell shortening and heart contraction is stored^5^. Ca^2+^ influx from the voltage-activated cardiac Ca^2+^ channel, Ca_V_1.2, concentrated in T-tubules, triggers release of SR Ca^2+^ from Ryanodine receptor-2 (Ryr2), located in the SR membrane, only 12 nm from the T-tubule. However, multiple other features of HF pathobiology localize to this critical cellular domain. There is a reduction in channel conductance, with fewer pore-forming subunits of Ca_V_1.2 channels ^6–8^. The remaining channels are more likely to be encoded by different splice variants and have increased open probability^9–11^. In HF, there is also spontaneous release of Ca^2+^ from the SR due to phosphorylation or oxidation of Ryr2, which is an important cause of delayed after-depolarization and arrhythmia^12^. This continuous Ryr2 leak depletes SR stores, resulting in smaller transients and less forceful contractions^12^. Depletion of SR Ca^2+^ is exacerbated by the reduced activity of the Ca^2+^ ATPase, SERCA2, in HF^13,14^. There are conflicting reports on whether or not SERCA levels change in HF^13,15,16^, though there is general consensus that its function is reduced as a consequence of decreased phosphorylation of Phospholamban (Pln)^17–19^. SERCA and therefore cell relaxation are inhibited by Pln in its unphosphorylated state; Pln inhibition of SERCA is relieved by Protein Kinase A (PKA) and Ca^2+^/calmodulin-dependent protein kinase II (CaMKII) phosphorylation and is maintained by protein phosphatase 1^18^.

Proximity proteomics is a technique well-suited to studying changes in the membrane-bounded space, known as the cardiac dyad, which regulates Ca^2+^-induced Ca^2+^-release and cardiac contractility^5,20^. Biotin-phenoxyl radicals generated upon H_2_O_2_ activation of the engineered plant peroxidase, APEX2, are unable to cross cell membranes^21^. Additionally, the short half-life of these radicals in the cytosol (∼1 ms) ensures that primarily proteins within close proximity (∼15-20 nm) are covalently modified. The high catalytic activity of APEX2 is sufficient to capture cell signaling with impressive temporal resolution (seconds to minutes)^22^. Indeed, cardiomyocytes from transgenic mice with fusion of APEX2 to the Ca_V_1.2 α_1C-_ and β_2B_-subunits were used to show that within minutes of exposure to a β-adrenergic agonist, biotin labeling of the RGK family protein Rad is significantly diminished, indicating decreased abundance and/or proximity of the protein to Ca_V_1.2 in the dyad^23^. Subsequent work indicated that in a manner similar to adrenergic activation of SERCA by Pln phosphorylation, PKA phosphorylation of Rad dissociates it from the membrane, relieving its inhibition of Ca_V_1.2 and generating greater Ca^2+^ influx and stronger contractions^20,23,24^. Using these transgenic mice, here we examine the proteome of the cardiac dyad in a mouse model of chronic ischemic cardiomyopathy. In mice with heart failure, we find enrichment of Galectin-1 (Gal-1), a lectin-binding protein, in the cardiac dyad.

Gal-1 is a ubiquitous, dimer-forming protein expressed in the cytoplasm and secreted, where it crosslinks glycosylated proteins^25^. It has anti-inflammatory effects, with implications for tumor biology and resolving inflammation after myocardial infarction (MI). Gal-1 expression is increased in the hearts of patients with ischemic and non-ischemic HF^26^. Global Gal-1 knockout (KO) mice have mildly reduced systolic function and worse remodeling post-MI, which was attributed to immune cell infiltration rather than through any effects in cardiomyocytes^26^. How the deletion of Gal-1 impairs systolic function is not known. There are reports that Gal-1 interacts with Ca_V_1.2^27^, though as an inhibitor. This feature would not explain the reduced systolic function in Gal-1 KO mice, though may, to some degree, explain the reduction in Ca^2+^ influx in systolic HF. As such, the enrichment of Gal-1 in the cardiac dyad in HF we observed suggests a potential role in HF pathobiology. In an effort to recapitulate the abnormal Ca^2+^ homeostasis seen in HF, and to clearly define the direct effects of Gal-1 on myocytes, we generated a mouse with over-expression of Gal-1 (Gal OE mice) in the heart. Rather than recapitulating the changes in HF myocytes, however, Gal OE mice have increased Ca_V_1.2 current and increased SERCA activity. Through generation of mice with over-expression of Gal-1, we show that Gal-1, long thought to exert its activity from outside of cells, is not merely present in the cardiac dyad, but activates both Ca_V_1.2 and SERCA.

## METHODS

### Data Availability

The data underlying this article will be shared on reasonable request to the corresponding author. The mass spectrometry proteomics data have been deposited to the ProteomeXchange Consortium via the PRIDE partner repository with the dataset identifier PXD063582 and 10.6019/PXD063582^28,29^.

### Animals

The Institutional Animal Care and Use Committee at Columbia University Vagelos College of Physicians and Surgeons approved the use of animals and the study protocol. Mice were fed a chow diet and housed in a barrier facility with 12 hour /12 hour light/dark cycles. Male and female mice of mixed background, aged 2 to 6 months, were used in the study. Experiments were conducted in a blinded fashion with respect to genotype and treatment.

### Transgenic APEX2-α_1C_ and APEX2-β_2B_ mice

Transgenic APEX2-α_1C_ and APEX2-β_2B_ mice were mated with cardiac-specific (α-MHC), doxycycline-regulated, codon-optimized reverse transcriptional trans-activator (rtTA) mice to produce double transgenic mice^30^. Mice were given ad libitum access to chow impregnated with doxycycline at 200 mg/g of chow (Bio-Serv S3888) 24 hours prior to euthanasia.

### Gal OE mice

The coding sequence for mouse Galectin-1 (Gal-1; GenBank: X66532.1), fused at the C-terminus with a linker and 3XFLAG tag, was inserted into the CTV targeting vector (Addgene #15912) downstream of the CAG promoter and a lox-stop-lox cassette by the Herbert Irving Comprehensive Cancer Center Genetically Modified Mouse Model Shared Resource at Columbia University. This targeting vector was recombined into the Rosa26 locus of the KV1 129-B6 hybrid embryonic stem (ES) cell line to generate targeted ES clones. These clones were injected into C57BL/6J blastocysts and implanted into surrogate mothers to produce chimeric mice. Offspring exhibiting germline transmission were crossed with Myh6 MerCreMer mice (Jackson Laboratory strain #005657) for cardiac-specific expression.

At five weeks, Gal-1 overexpression was induced via intraperitoneal (i.p.) injection of tamoxifen (40 mg/kg in soybean oil) every 48 hours for a total of five doses, following a protocol designed to minimize Cre- and tamoxifen-associated toxicity^31^.

### APEX2 proximity labeling

For proteomics experiments, hearts from mice with transgenic expression of Ca_V_1.2 APEX2-α_1C_ and APEX2-β_2B_ were rapidly removed and placed in ice-cold PBS. The atria were removed. For failing hearts with myocardial infarction, the scar and a generous margin of surrounding tissue were excised. The remaining heart tissue was chopped into 2-3 mm pieces. Preliminary studies in the laboratory showed that biotin labeling can reach a depth of ∼1 mm in intact tissue. Heart pieces were incubated with room temperature PBS or biotin-phenol 0.5 M for 30 minutes. For each heart, one aliquot of biotin-phenol treated tissue was then incubated with 1 M H_2_O_2_ for one minute to initiate biotinylation of nearby proteins. All samples were then treated with ice cold quenching solution containing (in mM) 10 sodium ascorbate (VWR 95035-692), 5 Trolox (Sigma 238813), and 10 sodium azide (Sigma S2002)^23^. Heart pieces were frozen at -80 C and subsequently homogenized with a rotostator (Omni) in lysis buffer containing (in mM), 50 Tris (tris(hydroxymethyl)aminomethane), 150 NaCl, 10 EGTA, 10 EDTA, 1% Triton X-100 (v/v), 0.1% SDS (w/v). Successful biotin labeling was confirmed for each heart by blotting with horseradish (HRP) peroxidase conjugated to streptavidin (Thermofisher, S911) at 0.3 μg/mL. Lysates containing 1 mg of protein were affinity purified and prepared for mass spectrometry as previously described^23^. Significant changes in protein enrichment were defined by a P value less than 0.05. Gene ontology enrichment analysis was performed using ToppGene (https://toppgene.cchmc.org/enrichment.jsp).

For imaging experiments, isolated ventricular cardiomyocytes from transgenic APEX2-α_1C_ mice with and without heart failure were plated on laminin (Millipore Sigma, L2020) coated coverslips.

Cells were incubated in PBS with 0.5 μM biotin-phenol (Iris-biotech) for 30 minutes, followed by 1 M H_2_O_2_ for one minute, prior to three 5 minute washes with ice-cold quenching solution. Cells were subsequently fixed in 4% paraformaldehyde. After blocking with 3% bovine serum albumin (MP Biomedicals, 9048-46-8), cells were incubated overnight with streptavidin Alexa Fluor™ 488 (Thermo Fisher, S11223) at 1.3μg/mL and imaged with confocal microscopy.

### Galectin-1 dimer structure predictions

Because Gal-1 is reported to dimerize through its C-terminus, and recombinant Gal-1 has C-terminal fusion of a linker and 3XFLAG motif, we modeled recombinant Galectin-1 dimerization with AlphaFold Multimer. Predictions were run on the San Diego Supercomputer Center cluster and accessed through the Cryo-EM Open Source Multiplatform Infrastructure for Cloud Computing (COSMIC2) portal^32,33^. Interface predicted modeling scores for these predictions were generated from AlphaFold3^34^. Native and recombinant Galectin-1 heterodimers were derived from the coding sequence of mouse Galectin-1 (X66532.1) and the coding sequence of Galectin-1 3XFLAG. Recombinant Galectin-1 homodimer predictions were derived exclusively from the coding sequence of Galectin-1 3XFLAG.

### Immunoblotting

Isolated cardiomyocytes were stored at -80 °C prior to homogenization in the lysis buffer described above, with the addition of 10 μM Calpain inhibitor I (Sigma A6185), 10 μM Calpain inhibitor II (Sigma A6060), and cOmplete Mini-tablets (Sigma 11836170001). For studies using phospho-specific antibodies, the phosphatase inhibitor tablet PhosSTOP (Roche) was dissolved in the buffer. Proteins were size fractionated with SDS PAGE, transferred to nitrocellulose (Bio-Rad 1620168 and Sigma GE10600012) and probed with the specified antibody. Antibodies are listed in **Supplemental Table 1**. Chemiluminescence signal was captured on an Azure Biosystem 600 imager and signal densitometry was normalized to TotalStainQ fluorescence loading control (Azure AC2227) and quantified using ImageJ. Ryr2 blots were imaged and quantified in image studio (LI-COR Biosciences).

### Whole-cell patch clamp electrophysiology studies

Mouse ventricular myocytes were isolated by enzymatic digestion as previously described^11^. Cardiomyocytes were placed in Bioptechs Delta T dishes containing 112 mM NaCl, 5.4 mM KCl, 1.7 mM NaH_2_PO_4_, 1.6 mM MgCl_2_, 20.4 mM HEPES pH 7.2, 30 mM taurine, 2 mM DL-carnitine, 2.3 mM creatine and 5.4 mM glucose. The dishes were mounted on the stage of an inverted microscope and served as a perfusion chamber. The pipette resistance was between 0.5 – 1.5 MΩ. The pipette solution contained 40 mM CsCl, 80 mM cesium gluconate, 10 mM 1,2-bis(o-aminophenoxy)ethane-N,N,N′,N′-tetraacetic acid, 1 mM MgCl_2_, 4 mM Mg-ATP, and 10 mM 4-(2-hydroxyethyl)-1-piperazineethanesulfonic acid (HEPES), adjusted to pH 7.2 with CsOH. After cardiomyocytes were dialyzed and buffered with 10 mM BAPTA in the internal solution, cells were locally superfused with 140 mM tetraethylammonium chloride, 0.5 mM BaCl_2_, 1 mM MgCl_2_, 5 mM glucose and 10 mM HEPES, adjusted to pH 7.4 with CsOH. A ramp protocol with a 200-ms voltage ramp from −60 mV to +30 mV was applied every 3 seconds. Voltage was corrected for liquid junction potential (−10 mV). Leak currents were subtracted by a P/3 protocol. Capacitance transients and series resistance were compensated (>88%). Under these conditions, currents through Ca^2+^ channels showed practically no inactivation. After establishing stable records, control traces were recorded. After the response to 100 nM isoproterenol superfusion stabilized, additional traces were recorded. The ramp protocol experiment analysis was previously described^11^.

### Cell contractility and transient analysis

Isolated cardiomyocytes were allowed to settle on laminin-coated coverslips. Cells underwent a stepwise Ca^2+^ tolerization protocol with CaCl_2_ to a final concentration of 1.2 mM in Tyrodes solution containing: 134 mM NaCl, 5.4 mM KCl, 1.2 mM CaCl_2_, 1 mM MgCl_2_, 10 mM HEPES, and 10 mM glucose. Myocytes were field-stimulated at 1 Hz. In the contractility study, recordings were made using an Ionoptix system. After contraction was stable for 30 seconds, cells were treated with rapid perfusion of 200 nM isoproterenol. Parameters of contraction were generated through monotonic analysis of the average of at least 8 contractions in the SarcLen module of IonWizard software (Ionoptix).

In the Ca^2+^ fluorescence study, cells were treated with 2.5 mM FURA2-AM for 20 minutes in Tyrode’s solution. After 2 washes, cells on coverslips were imaged in a perfusion as described^20^. Cells were paced for 1 minute prior to recording Ca^2+^ transients for 10 seconds. Similarly, after 2 minutes of perfusion with 100nM isoproterenol, responses were recorded for 10 seconds. SR Ca^2+^ load was determined with perfusion of 10 mM caffeine. All experiments were recorded in Nikon Elements and analyzed in MATLAB^20^. The NCX rate constant was calculated from the decay of SR Ca^2+^ release after rapid application of caffeine (1/τ_caffeine_), from which the SERCA2 constant was also calculated (1/τ_1Hz_-1/τ_caffeine_)^13^.

### Quantitative Real-Time Reverse Transcription Polymerase Chain Reaction

RNA was extracted from isolated cardiac myocytes using the Quick-RNA Miniprep Plus (Zymo Research). cDNA was synthesized from total RNA using High-Capacity cDNA Reverse Transcription Kit (Applied Bio Systems). The qPCR was performed using SYBR mix (Thermo Fisher Scientific) on a StepOnePlus Real Time PCR system (Applied Bio Systems). Transcript quantification for mRNAs was performed using the Δ-Δ method using primers designed for Cacna1c, the gene encoding Ca_V_1.2, and 18S, the internal control. Primer sequences used for independent validation:

Cacna1c:

Forward: GCTTATGGGGCTTTCTTGCAC

Reverse: ACTGGACTGGATGCCAAAGG

18S:

Forward: GTAACCCGTTGAACCCCATT

Reverse: CCATCCAATCGGTAGTAGCG

### Echocardiography

8-to 16-week-old mice were anesthetized with 1% to 2% isoflurane and placed on a heated table in the supine position. Transthoracic echocardiography was performed at the Columbia University Oncology Precision Therapeutics and Imaging Core using the Vevo 3100 Imaging System (FUJIFILM VisualSonics). Vevo LAB 3.0 analysis software was used to measure chamber dimensions. M-mode images were used to quantify left ventricular diameters and ejection fraction (EF%). Vevo Strain Software was used to calculate left ventricular volume and mass in Gal OE mouse experiments and, for speckle-tracking strain analysis, determination of global circumferential strain in all mice. Pulse wave doppler recordings of mitral inflow were obtained in the apical four-chamber view, as were tissue doppler recordings of the lateral mitral annulus.

### Electrocardiography

Electrocardiograms were recorded on isoflurane-anesthetized mice 8 to 12 weeks of age using an Emka system with subcutaneous electrodes and IOX software (Emka Technologies). In the adrenergic stimulation study, after recording baseline electrogram for 2 minutes, mice were given isoprotenol 2 mg/kg i.p. followed by recording for 5 minutes. Three randomly selected measurements of PR, RR, QRS, and QT duration were made using IOX software and averaged. The QT duration was corrected for heart rate using Mitchells’s formula^35^.

### Telemetry

8 to 12 week-old mice were housed singly after subcutaneous implantation of radiotelemeters (DSI, ETA-F10). After one week of recovery, the cage was placed on a radio receiver and a baseline electrocardiogram was recorded for 1 hour. Subsequently, mice were injected with epinephrine 2 mg/kg, and recordings were continued for 23 hours. The presence of arrhythmia was assessed using Ponemah software with the Data Insights rhythm analysis module.

## RESULTS

### Generating systolic heart failure in Ca_V_1.2 APEX2-α_1C_ and APEX2-β_2B_ transgenic mice

We have previously shown that myocytes from transgenic mice with cardiac-specific inducible expression of Ca_V_1.2 APEX2-α_1C_ and APEX2-β_2B_ have normal contractile function and responses to β-adrenergic stimulation^23^. Expression of transgenic APEX2-α_1C_ and APEX2-β_2B_ is regulated by a TetO element and mice for both strains are crossed with MHC-rtTA mice to drive expression upon providing mice doxycycline impregnated chow (**Figure 1A**). Heart failure was generated at 8 to 12 weeks of age through surgical ligation of the left anterior descending coronary artery (**Figure 1B**). In order to study the effect of chronic remodeling of the infarcted heart that develops during ischemic cardiomyopathy, m-mode, 2-D and strain echocardiography were performed on surviving mice starting 2-months after myocardial infarction and serially until severe heart failure was present. (**Figure 1B**). Mice with myocardial infarction had significantly decreased systolic function assessed by fractional shortening (FS), fractional area of change (FAC), and global circumferential strain (GCS), compared with healthy control (Ctrl) APEX2-α_1C_ and APEX2-β_2B_ mice littermates (**Figure 1C-D** and **S1B**). Mice in the infarcted group also had significantly dilated ventricles, assessed as end-diastolic volume (EDV) and left ventricular end-diastolic diameter (LVEDD), consistent with chronic remodeling of the left ventricle (**Figure 1E-F** and **S1A**). Other signs of HF in the coronary ligation group include wall thinning, aneurysm formation, increased heart weights, and increased lung weights (**Figure S1C-E**).

**Figure 1.**
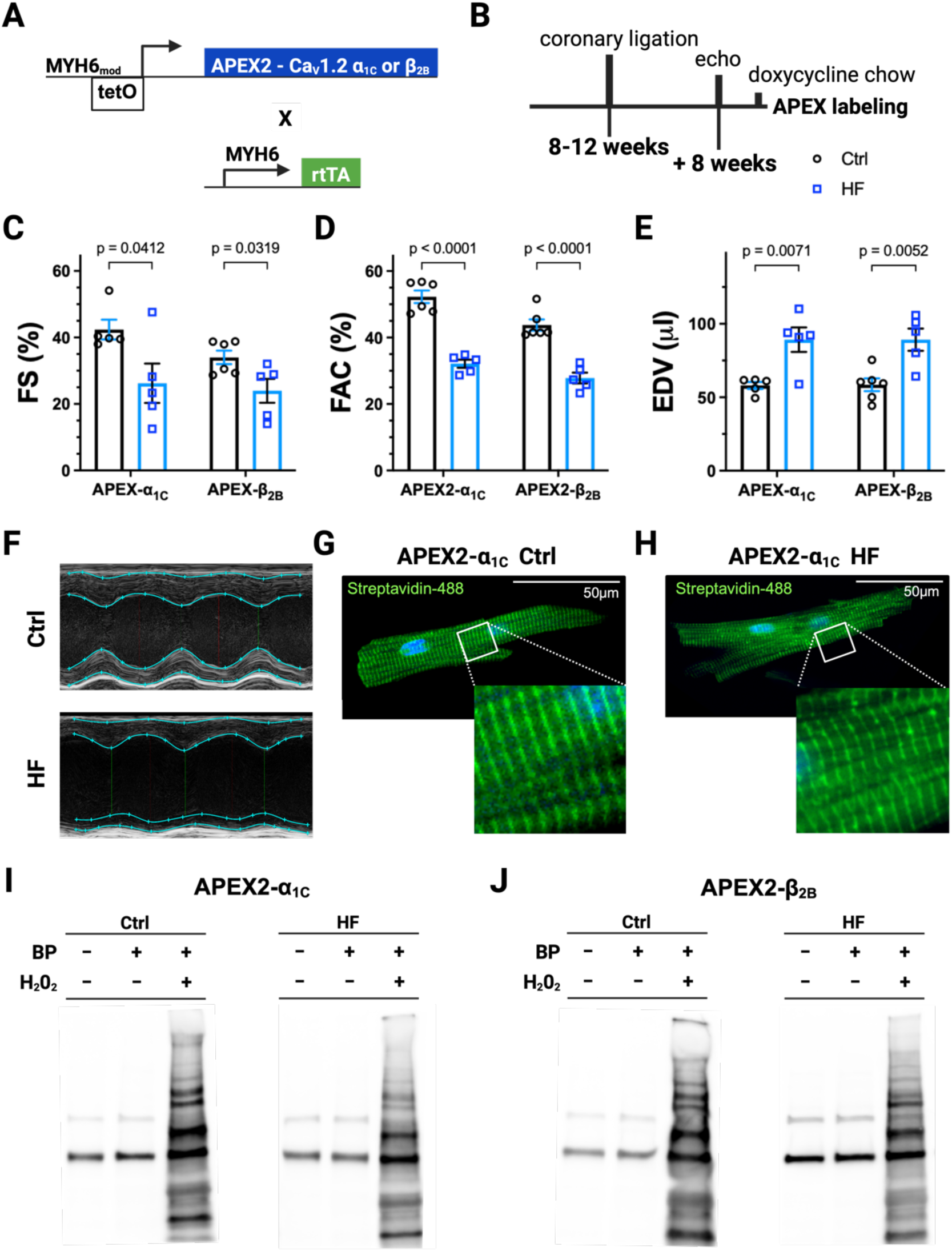
Generating HF in APEX2-α_1C_ and the APEX2-β_2B_ mice and proximity labeling the Ca_V_1.2 interactome. **A**. Schematic showing the design of tetracycline inducible APEX2-α_1C_ and APEX2-β_2B_ mice. **B**. Experimental timeline for expression of transgenic Ca_V_1.2 subnits and generation of HF. **C**, **D**, and **E**. Fractional shortening, (FS), fractional area of change (FAC), and end diastolic volume (EDV) for transgenic mice 8 weeks after coronary artery ligation (HF) and control (Ctrl). **F**. Representative M-mode images o APEX2-β_2B_ mice without (upper panel) and with HF (lower panel). **G** and **H**. Representative immunofluorescence images of myocytes isolated from APEX2-α_1C_ mice showing the localization of APEX2 catalyzed biotinylation of Ca_V_1.2 neighbors after treatment with biotin-phenol and H_2_O_2_, prior to fixation and treatment with streptavidin conjugated to Alexa Fluor 488 and DAPI. **I** and **J**. Streptavidin-HRP blots of heart tissue lysates from APEX2-α_1C_ and APEX2-β_2B_ mice with and without HF and treated with and without biotin-phenol and H_2_O_2_. For echo studies, N=6 mice for Ctrl and N=5 mice for HF, for both APEX2-α_1C_ and APEX2-β_2B_ mice. Indicated P-values were calculated by unpaired two-tailed Student’s T-test.

### Defining the cardiac dyad proteome of systolic heart failure in mice

Left ventricular myocytes were isolated after confirmation of significant systolic heart failure by echocardiography, defined as FAC less than 35%. The dyad proteome was labeled in cardiomyocytes from healthy mice and mice with HF after incubation in biotin-phenol for 30 minutes, followed by APEX2 activation with H_2_O_2_. Correct localization of transgenic channels and successful proximity labeling were noted in cells from transgenic APEX2-α_1C_ mice in both groups by confocal microscopy (**Figure 1G-H**). Successful biotin labeling was also confirmed for each mouse included as a replicate in the mass spectrometry experiment. For both APEX2-α_1C_ and APEX2-β_2B_ mice, 6 Ctrl and 5 mice with HF, defined here by FAC less than 35%, were studied. Mice with and without HF demonstrated similar and robust levels of biotin-labeling upon streptavidin blotting heart lysates (**Figure 1I-J**). Following affinity purification of heart lysates for biotinylated proteins, TMT labeling, and triple-stage mass spectrometry, we quantified 2326 proteins in APEX2-α_1C_ mice hearts, of which 71 were significantly altered in HF (**Figure 2A, Data Supplement**). In APEX2-β_2B_ mice hearts, we quantified 2243 proteins, of which 336 were significantly altered in HF (**Figure 2B, Data Supplement**).

**Figure 2.**
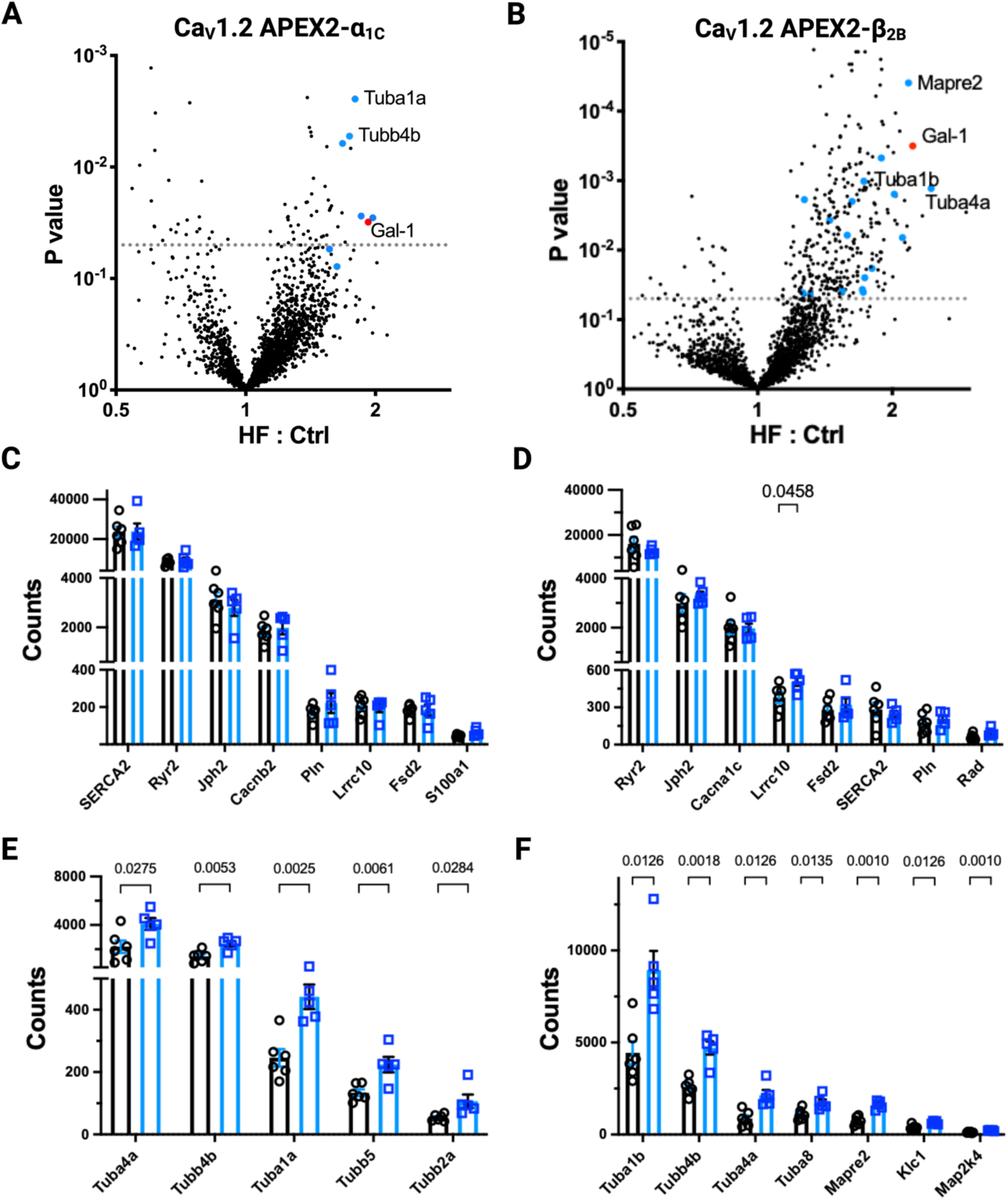
Mapping the dyad interactome in HF with APEX2-Ca_V_1.2 mice. **A** and **B**. Volcano plots of fold-chage in protein quantification by TMT mass spectrometry after streptavidin-based affinity purification for APEX2-α_1C_ and the APEX2-β_2B_ mice with and without HF. Microtubule proteins are colored blue. Galectin-1 is colored red. Dotted line indicates a P-value of 0.05. **C** and **D**. Relative differences in quantification of dyad-specific proteins. Indicated P-values are included for proteins significantly different in mice with HF. **E** and **F**. Quantification of microtubule specific proteins enriched in the interactome of APEX2-α_1C_ and APEX2-β_2B_ mice with HF. For all experiments, P-values below the threshold for significance (p = 0.05) are shown. Indicated P-values are included for proteins significantly different in mice with HF. N=6 mice for Ctrl and N=5 mice for HF, for both APEX2-α_1C_ and the APEX2-β_2B_ mice. Indicated P-values were calculated by two-tailed T-test.

### Dyadic remodeling in HF does not affect the relative proximity and abundance of Ca^2+^ -handling proteins

In the interactome of transgenic Ca_V_1.2 mice with HF, we found no difference in the enrichment of Ryr2, Pln, SERCA2, or several other proteins described as localizing to the cardiac dyad (**Figure 2C-D**)^36^. One notable exception, however, was the 36% increase in the leucine-rich repeat containing protein 10 (Lrrc10) in the interactome of β_2B_-APEX2 mice with HF (p = 0.0458). Lrrc10 forms complexes with Ca_V_1.2, increases channel activation, helps maintain heart function after MI and transverse aortic constriction, and has been leveraged in peptide-based strategies to increase Ca_V_1.2 currents^37–39^. Compared to controls, mice with HF in both transgenic lines demonstrate significant enrichment of microtubule proteins in the vicinity of the channel (**Figure 2E-F**). This is consistent with reports showing marked expansion of sub-sarcolemmal microtubules throughout the myocyte in HF as well as reports that microtubule mediated delivery of ribosomes and mRNA to the periphery is a requirement in the development of myocyte hypertrophy^40,41^.

### Identification of the ubiquitin-proteasome, translation machinery, and Gal-1 in the dyad in HF

Gene ontology (GO) enrichment analysis for both transgenic mice showed significant enrichment of the ubiquitin-proteasome system (**Figure 3A-D**). The total amount of the 26S proteasome non-ATPase regulatory subunit 4 (Psmd4) was compared in lysates from APEX2-α_1C_ and APEX2-β_2B_ mice with and without HF, without streptavidin affinity purification (**Figure 3E-F**). There was no difference in total Psmd4 levels in HF compared to Ctrl hearts (p = 0.7381 and p = 0.669 for APEX2-α_1C_ and APEX2-β_2B_ mice, respectively) (**Figure 3G**). This suggests that the significant increase in biotinylated Psmd4 in HF hearts reflects closer interaction of the channel to the proteasome in HF, to the exclusion of increased abundance.

**Figure 3.**
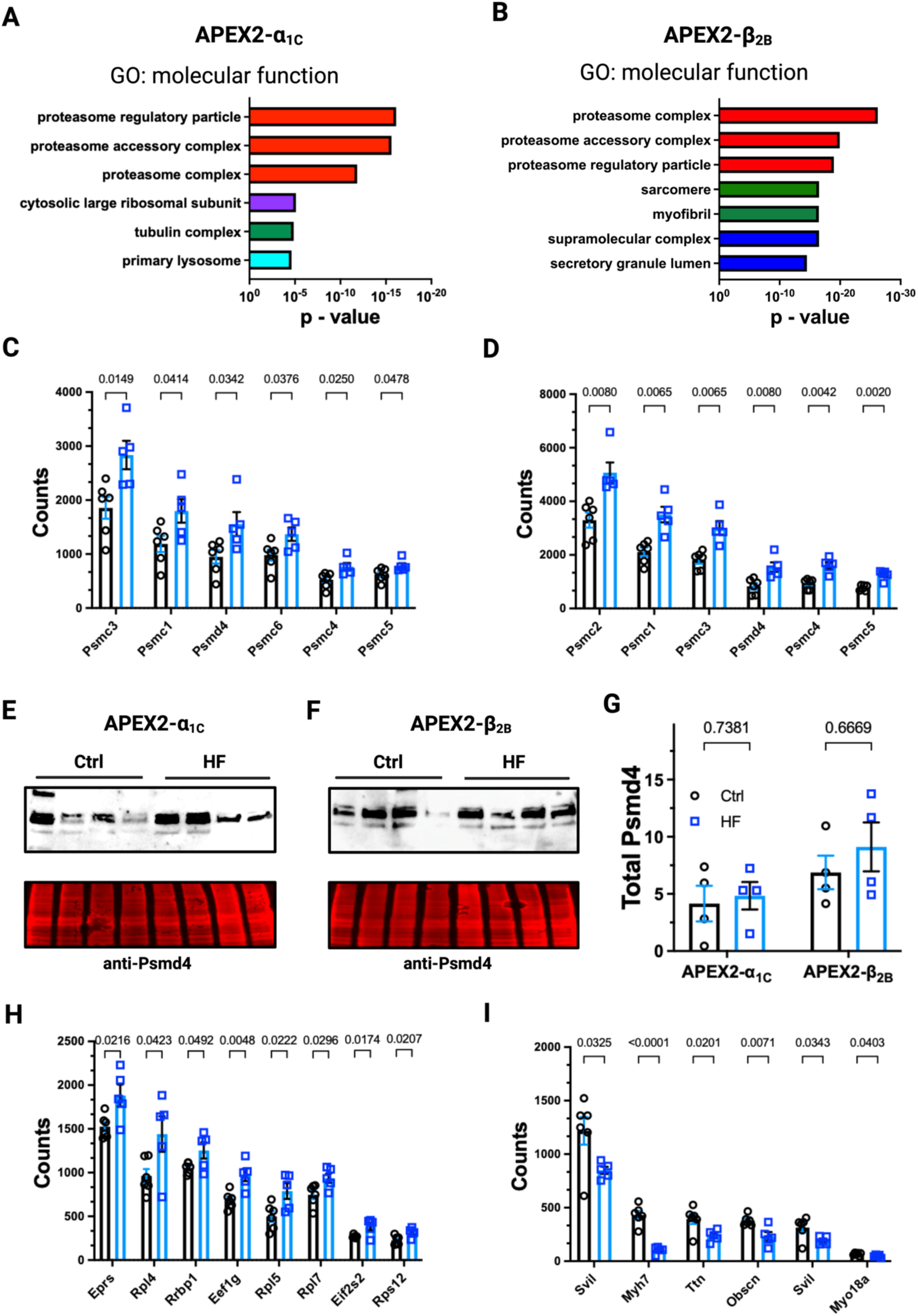
Heart failure induced changes in the cardiac dyad includes enrichment of the ubiquitin-proteasome. **A** and **B**. Gene ontology analysis of proteins significantly enriched in the proximity proteome of APEX2-α_1C_ mice and APEX2-β_2B_ mice with HF. **C** and **D**. Relative differences in enrichment of ubiquitin-proteasome proteins in the Ctrl and HF APEX2-α_1C_ and APEX2-β_2B_ mouse heart interactome. **E** and **F**. Anti-Psmd4 immunoblot of Ctrl and HF APEX2-α_1C_ and APEX2-β_2B_ mouse heart (upper panels), with protein loading controls (lower panels). **G**. Quantification of anti-Psmd4 signal densitometry is normalized to protein-loading controls. Each replicate is derived from a single mouse. N=4 for WT and N=4 for both APEX2-α_1C_ and APEX2-β_2B_ mice. **H**. Relative differences in enrichment of moyfilament proteins in interactome of Ctrl and HF APEX2-α_1C_ mouse heart. **I**. Relative differences in quantification of proteins involved in protein translation in Ctrl and HF APEX2-α_1C_ heart. N=6 for WT and N=5 for HF. Indicated P-values were calculated by two-tailed T-test.

In mice with HF, the APEX2-α_1C_ interactome was also particularly enriched for proteins associated with protein translation including the ribosome (**Figure 3H**), while myofilament proteins were labeled less (**Figure 3I**). This might reflect the dyadic widening that develops with remodeling in HF and the increased distance from T-tubular APEX2-α_1C_ subunits to the underlying contractile machinery^2^. Gal-1 was significantly enriched in both HF groups: it increased 1.9-fold in the APEX2-α_1C_ dyad (p = 0.0312) and 2.2-fold in the β_2B_-APEX2 dyad (p = 0.001) (**Figure 2A-B**).

### Galectin-1 overexpression increases calcium current

Gal-1 can form homodimers through non-covalent interactions at the C-terminus, with a K_d_ of 7μM (**Figure 4A**)^42^. Addition of a linker and 3XFLAG to the C-terminus was not predicted to disrupt formation of heterodimers with native Gal-1 or homodimers of recombinant Gal-1 (**Figure 4A-B**). Gal-1 overexpressing mice (Gal OE) have Cre-inducible expression of Gal-1-3XFLAG and are crossed to cardiac myosin heavy chain mER-Cre-mER mice for cardiomyocyte-specific expression (**Figure 4C**). Gal OE progeny are bred in typical Mendelian ratios. Homozygous Gal OE and wildtype Cre-positive littermates were induced with tamoxifen beginning at 5 weeks, followed by a 2-week washout period to mitigate any transient Cre cardiotoxicity^31^ (**Figure 4D**). Isolated myocytes from Gal OE contain a 17 kDa band on anti-Gal-1 immunoblotting that is absent in the wildtype Cre-positive littermates (WT) (**Figure 4E-F**). We conclude there is no transcriptional down-regulation of Gal-1 in Gal OE as native Gal-1 levels are unchanged (**Figure 4G**). Altogether, Gal OE myocytes have a 3.5-fold increase in Gal-1 relative to WT (p = 0.0064) (**Figure 4H**).

**Figure 4.**
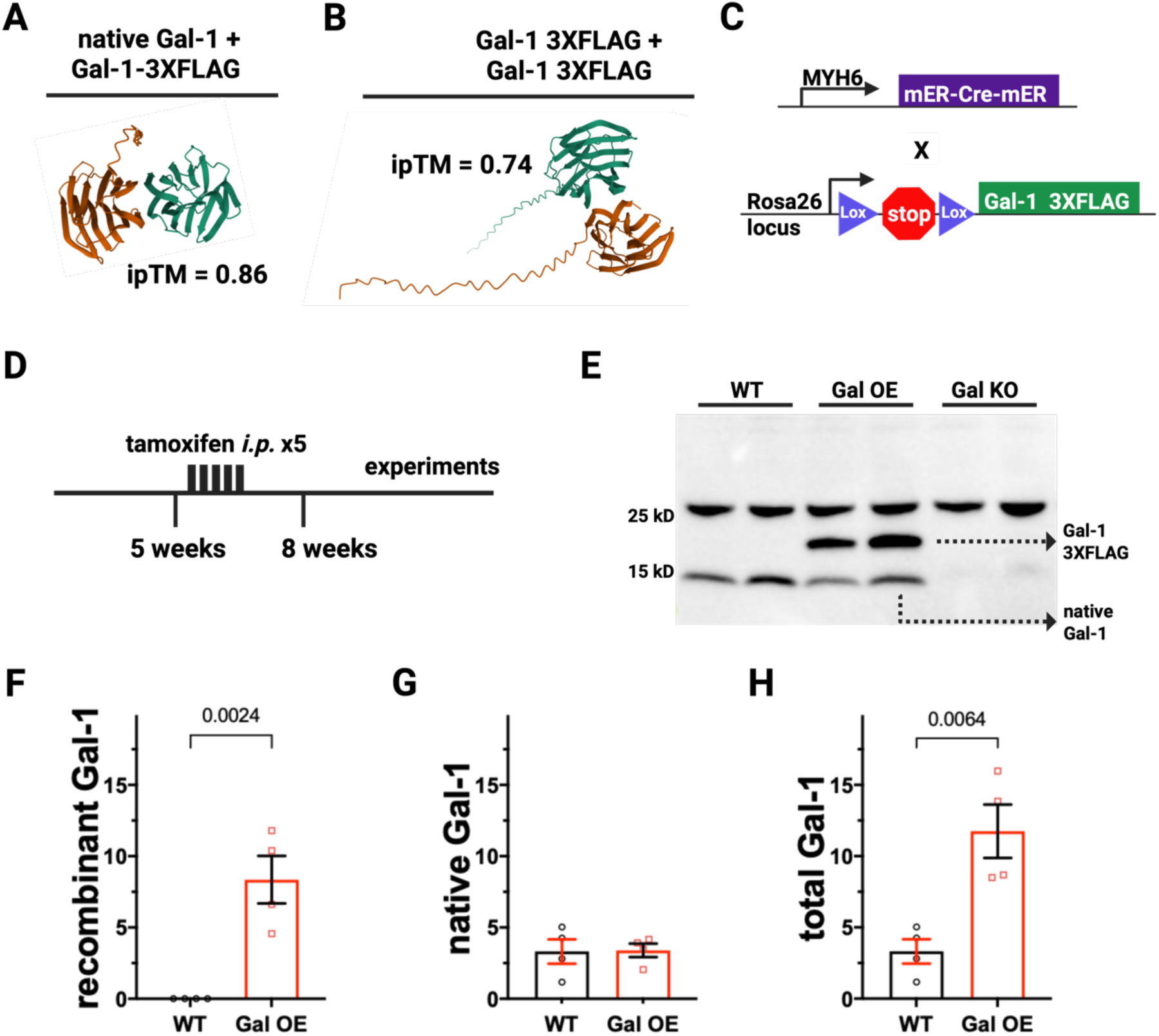
Galectin-1 overexpression mice have robust, inducible expression of recombinant Gal-1. **A**. Alphafold Multimer structure prediction for native and recombinant Gal-1 3XFLAG heterodimer formation. iPTM values were calculated in Alphafold3. **B**. Alphafold Multimer structure prediction for Gal-1 3XFLAG homodimer formation (right structure). **C**. Schematic showing the design of the Gal-1 overexpression (Gal OE) mouse. **D**. Timeline for the tamoxifen treatment and experimentation on Cre-positive Gal OE and WT control mice. **E**. Representative anti-Gal-1 immunoblot of cardiomyocytes isolated from WT, Gal OE and Galectin-1 knockout mice. **F**, **G**, and **H**. Quantification of 14kDa native Gal-, 17kDa recombinant Gal-1, and total Gal-1 in WT and Gal OE mice. N=4 for both WT and Gal OE.

### Galectin-1 overexpression increases Ca^2+^ current and contractility

Myocytes isolated from Gal OE have increased Ca_V_1.2 current with a 19.4% increase in maximal conductance compared to WT (p = 0.0074) (**Figure 5A-B**). Cells from both Gal OE and WT mice respond appropriately to treatment with a saturating dose of the adrenergic agonist, isoproterenol, with a left-shift in activation and no difference in maximal conductance (**Figure 5C-D**). To determine if Gal-1 overexpression increases whole-cell currents through increasing the number of channels, lysates from Gal OE and WT myocytes were immunoblotted. Anti-Ca_V_1.2 α_1C_ immunoblots show no difference in channel expression (**Figure 5E-F**). Nor is there any evidence of transcriptional upregulation, with no difference in Cacna1c transcripts (**Figure 5G**). Similarly, there was no change in expression of Ca_V_1.2 β-subunits detected with a pan-β-subunit antibody (**Figure 5E** and **5H**). Isolated cardiomyocytes were loaded with FURA2-AM for ratiometric measurement of intracellular Ca^2+^. Ca^2+^ transients were in Gal OE mice were 43.7% greater than in WT controls (p = 0.0001), and 16.4% greater after isoproterenol treatment (p = 0.0199) (**Figure 5I**).

**Figure 5.**
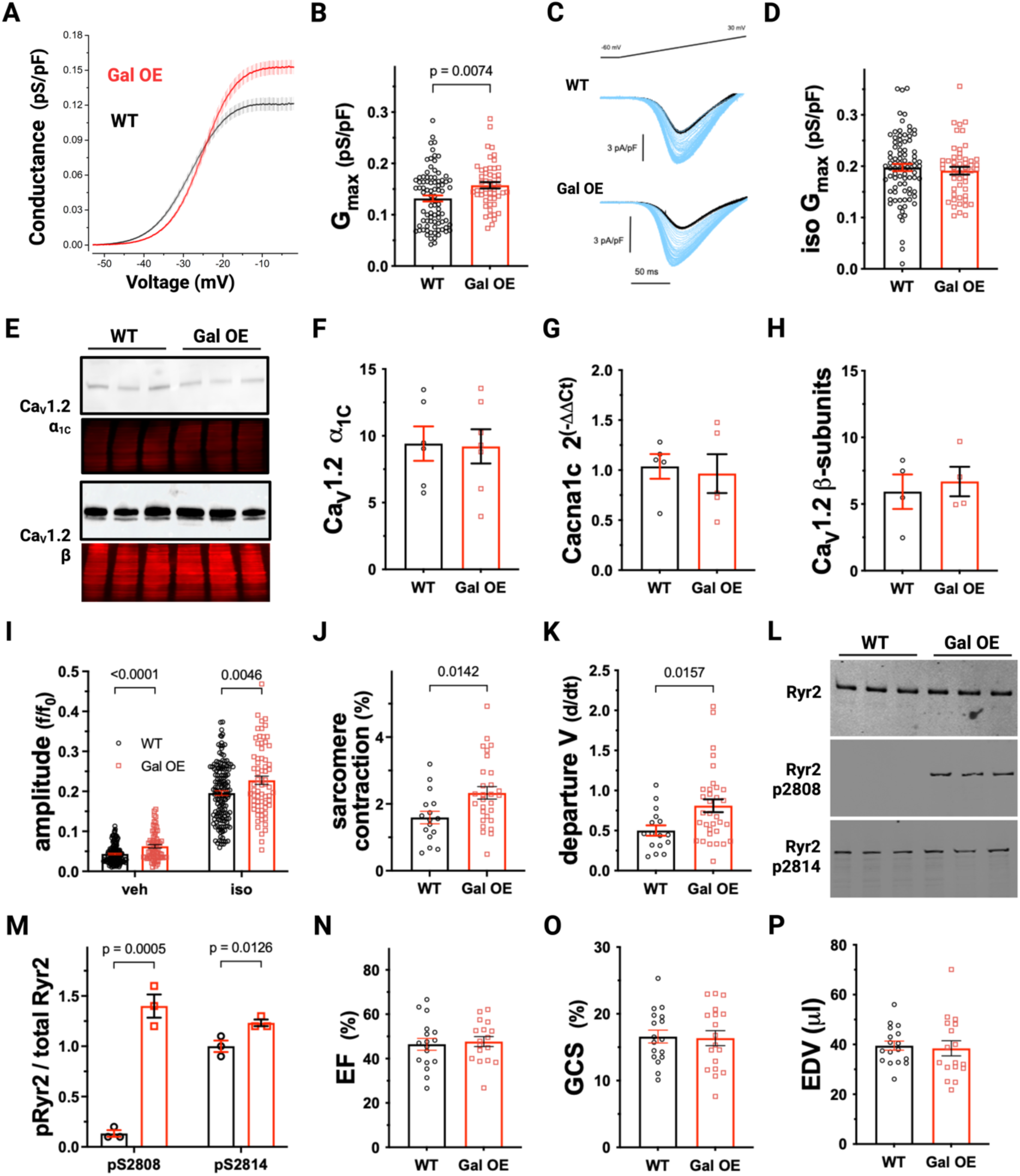
Gal OE mice have increased activation of Ca_V_1.2, increased Ca^2+^ transients and increased cell shortening. **A.** Plot of conductance density (G) and voltage for WT and Gal OE mice. Mean ± SEM. **B**. Maximal conductance density for Ca_V_1.2 currents in WT and Gal OE cardiomyocytes. **C**. Representative recordings of voltage-clamped WT and Gal OE cells subjected to 2 ms voltage ramps before (black trace) and after treatment with 100nM isoproterenol (blue traces). **D**. Maximal conductance density for isoproterenol treated WT and Gal OE cells. WT, N=7 mice, n=86 cells. Gal OE, N=4, n=51. Indicated P-values calculated by two-tailed Mann-Whitney T-test. **E**. Anti-Ca_v_α_1C_ and anti-Ca_v_β immunoblots of WT and Gal OE cardiomyocytes (upper panels), with protein loading controls (lower panels). **F**. Quantification anti-Ca_v_α_1C_ antibody signal densitometry, normalized to protein-loading controls in panel E. Each replicate is averaged data from a single mouse. N=6 for WT and N=7 for Gal OE. **G**. Quantitative PCR of Cacna1c transcripts from isolated WT and Gal OE myocytes. N=5 for WT and N=5 for Gal OE mice. **H**. Quantification of anti-Ca_v_β-subunit antibody signal, normalized to protein-loading controls in panel E. N=4 for WT and N=4 for Gal OE. Indicated P-values were calculated by two-tailed Student’s T-test. **I**. Amplitude of the Ca^2+^ transient was assessed in FURA2-AM treated WT and Gal OE myocytes paced at 1 Hz and perfused with vehicle (veh) or 100nM isoproterenol (iso). **J** and **K**. Sarcomere shortening and velocity of sarcomere shortening, or departure velocity (V), are increased in Gal OE cardiomyocytes. N=3 mice and n=16 for WT cells. N=6 mice and n=35 cells for Gal OE. **L**. Anti-Ryr2 immunoprecipitates from WT and Gal OE cardiomyocytes were blotted phosphorylated Ryr2^S2808^ and Ryr2^S2814^, both normalized to total Ryr2 signal. **M**. Quantification of phosphorylated Ryr2. N=3 for both WT and Gal OE. **N**. EF was calculated from parasternal long-axis views. **O**. GCS was calculated using VevoStrain software. **P**. EDV was calculated in VevoLab software from parasternal long-axis views. For analyses in panels N through P, N=17 for WT and N=18 for Gal OE. Statistical significance was determined by two-tailed Student’s T-test.

Isolated cardiomyocytes from Gal OE mice have increased sarcomere shortening (**Figure 5J**) and contract more quickly than cells from WT mice (**Figure 5K**). Concordant with the increase in Ca^2+^ transients and cell shortening, Gal OE myocytes have markedly increased phosphorylation of Ryr2^S2808^ (1.4 versus 0.1 for Gal OE and WT cells, respectively, p = .0005) and mildly increased phosphorylation of Ryr^S2814^ (1.2 versus 1.0 for Gal OE and WT cells, p = .0249), sites of PKA and CaMKII phosphorylation, respectively, that augment SR Ca^2+^ release (**Figure 5L-M**). To assess whether increased cell shortening in Gal OE cells results in increased contractile function at the organ level in healthy mice, we examined cardiac function with M-mode, 2D, and speckle-tracking echocardiography. Systolic function assessed by EF and GCS was unchanged (**Figure 5N-O**). Similarly, Gal-1 overexpression did not affect left ventricular volumes (**Figure 5P**).

### Galectin-1 overexpression increases lusitropy

Gal-1 overexpression resulted in accelerated Ca^2+^ removal during relaxation, with decreased τ, a parameter of transient decay, both before and after isoproterenol treatment (**Figure 6A**). Diastolic Ca^2+^ was unchanged by Gal-1 overexpression and decreased 0.8% in Gal OE mice after isoproterenol treatment. In WT mice, diastolic Ca^2+^ decreased 5.8%, to a significantly lower cytosolic concentration after isoproterenol (0.16 +/- 0.3 for Gal OE cells versus 0.15 +/- 0.2 for WT, p = 0.0016) (**Figure 6B**). SR Ca^2+^ load was assessed by rapid perfusion of caffeine, an agonist of Ryr2 Ca^2+^ release channels in the SR. Consistent with increased Ca_V_1.2 activation and faster removal of cytosolic Ca^2+^, higher Ca^2+^ peaks after caffeine treatment indicate Gal OE cardiomyocytes have larger SR Ca^2+^ stores (p = 0.0276) (**Figure 6C**). Rate constants for the Na^+^/Ca^2+^ exchanger (NCX) and SERCA2 were derived from caffeine treated cells as described by others^13^. The NCX rate constant was mildly increased and 12.7% faster for cells with Gal-1 overexpression (p = 0.0209) (**Figure 6D**), whereas the SERCA rate constant was 39.1% faster in Gal OE myocytes compared to WT (8.8 +/- 2.7 s-1 for Gal OE versus 6.3 +/- 3.5, p < 0.0001) (**Figure 6E**). Diastolic function was examined at the organ-level using doppler imaging of trans-mitral filling in healthy Gal OE and WT mice. The ratio of early-to-late left ventricular filling (E/A) was unchanged by Gal-1 overexpression in the heart (**Figure 6F**). Nor were there differences in E/e’ (**Figure 6G**) or isovolumetric relaxation time (IVRT) (**Figure 6H**), additional parameters of diastolic function. Immunoblots of SERCA2 and NCX1 indicate similar levels of these proteins (**Figure 7A-C**). Similarly, levels of Pln, the SERCA inhibitor, were unchanged by overexpression of Gal-1 (**Figure 7D-E**). Interestingly, increased phosphorylation of the Pln residues that relieve SERCA2 inhibition was detected in Gal OE cardiomyocytes (11.2 +/- 2.8 for Gal OE myocytes versus 5.7 +/- 1.8 for WT, p = 0.0151). (**Figure 7D** and **7F**).

**Figure 6.**
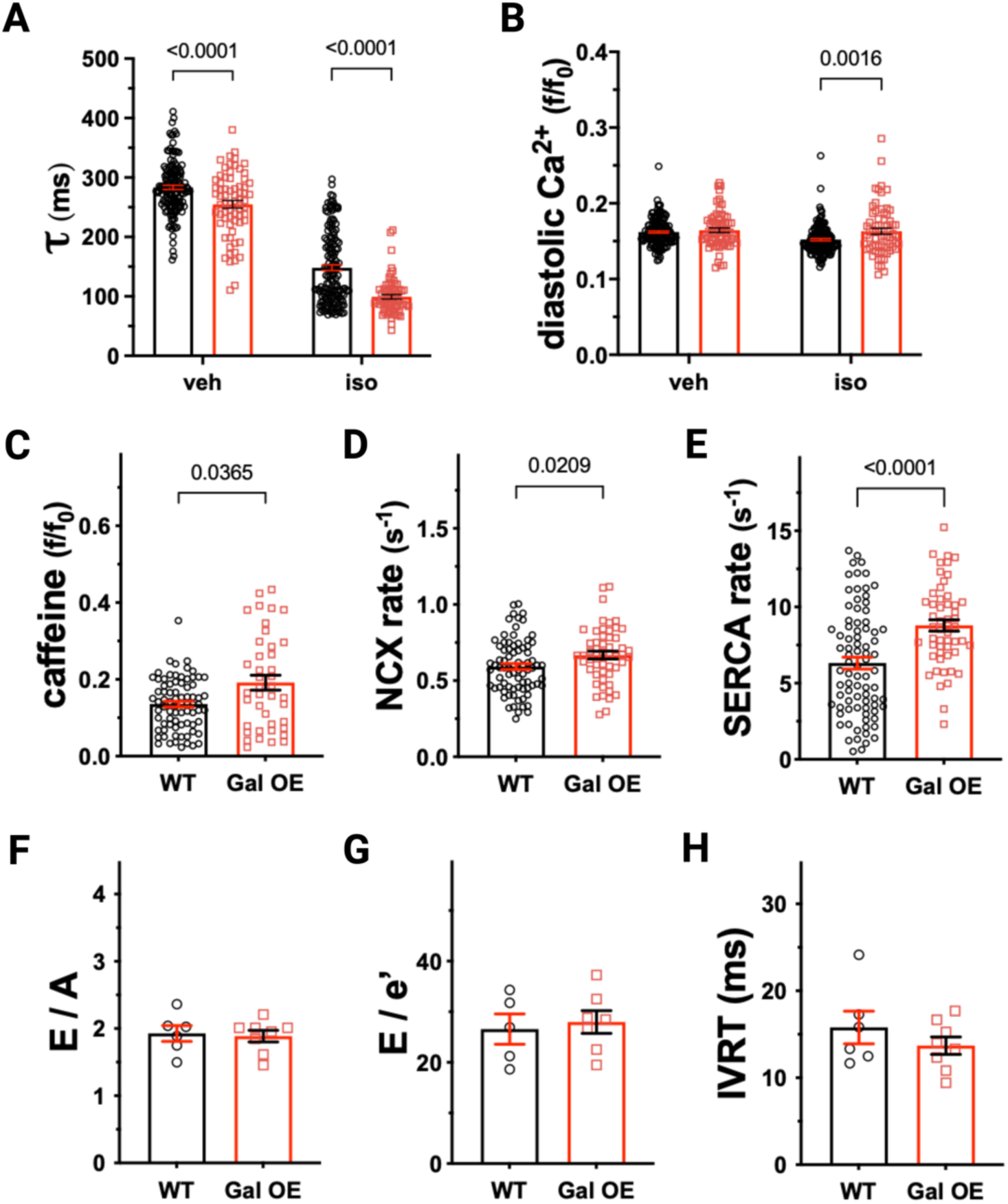
Gal-1 overexpression increases lusitropy. **A**. Ratiometric measurements of cytosolic Ca^2+^ in diastole were recorded for vehicle and isoproterenol treated cells. **B**. τ was calculated for transient decay in vehicle and isoproterenol treated cells. **C**. Ratiometric amplitude of Ca^2+^ peak after perfusion of 10 mM caffeine **D** and **E**. Rate constants for NCX1 and SERCA2 were calculated using decay of the caffeine Ca^2+^ peak (see methods). **F**. Left ventricular filling was characterized by as the average E/A ratio for each mouse. **G**. Diastolic function was also assessed as the average E/e’ obtained by mitral inflow doppler and tissue doppler of the lateral mitral annulus. **H**. Average isovolumetric relaxation time (IVRT) was compared for WT and Gal OE mice. For experiments in panels A and B, N=3 mice, n=154 cells for WT and N= 3 mice, n=69 cells for Gal OE. For panels C, E and E, N=3 mice, n=83 cells for WT. N= 3 mice, n=53 cells for Gal OE. Indicated P values were calculated using two-tailed Mann-Whitney T-test. For diastolic echocardiography studies (panels F, G, and H), N=6 for WT and N=8 for Gal OE. Significance was assessed by two-tailed T test.

**Figure 7.**
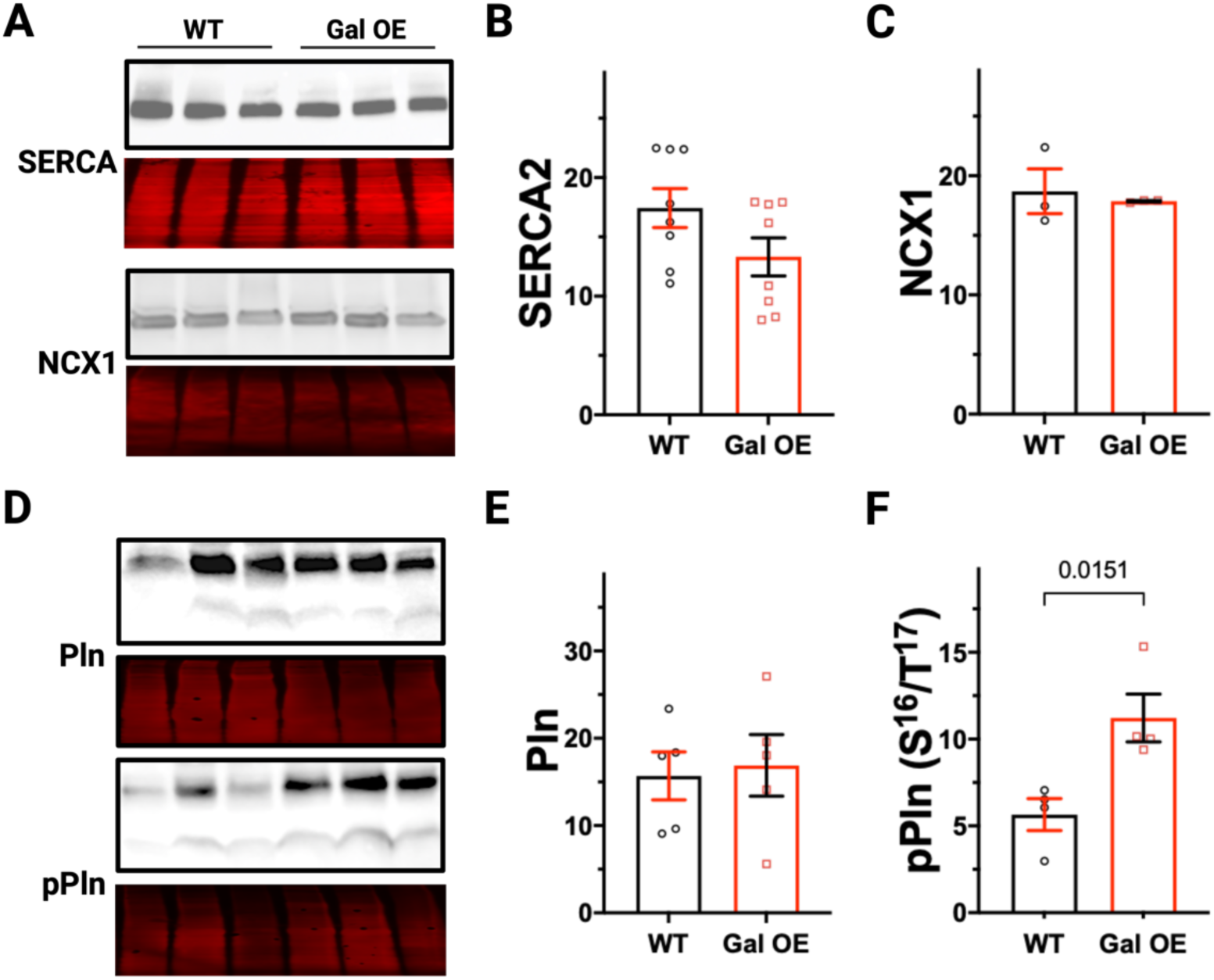
Phospholamban phosphorylation is increased in Gal OE hearts. **A**. Anti-SERCA2 and anti-NCX1 immunoblots of WT and Gal OE cardiomyocytes (upper panels), with protein loading controls (lower panels). **B** and **C**. Quantification of SERCA2 and NCX1 signal densitometry, normalized to protein-loading controls in panel A. Each replicate is averaged data from a single mouse. For SERCA2, N=8 for WT and N=8 for Gal OE. For NCX1, N=3 for WT and N=3 for Gal OE. **D**. Immunoblots with anti-Phospholamban (Pln) antibody and the phosphospecific anti-Pln S^16^/T^17^ (pPln) antibody (upper panels) and protein loading controls (lower panels). **E** and **F**. Quantification of Pln monomer signal, normalized to protein-loading controls in panel A. For Pln immunoblots, N=5 for WT and N=5 for Gal OE. For pPln, N=4 for WT and N=4 for Gal OE. Indicated P-values were calculated by two-tailed Student’s T-test.

### Gal-1 overexpression does not cause arrhythmia

In light of increased Ca^2+^ current in Gal OE myocytes, electrocardiographic parameters were examined. Mice were sedated with isoflurane anesthesia for recording of surface electrocardiograms. Heart rate was significantly lower in Gal OE mice than WT controls (496 +/- 8.9 beats per minute for Gal OE versus 527 +/- 35 for WT, p = 0.0377) (**Figure 8A**). PR interval, QRS duration, and corrected QT interval (QTc) were unchanged by Gal-1 overexpression (**Figure 8B-D**). As Gal OE myocytes have increased NCX and increased Ryr2 phosphorylation, potential precipitants of early and delayed after-depolarizations^43,44^, we recorded electrocardiograms for 5 minutes after i.p. injection of mice with isoproterenol. No arrhythmias were noted and Gal OE and WT control mice had similar, appropriate changes in HR, atrioventricular conduction, and QTc (**Figure 8E-G**). In case β-adrenergic initiated arrhythmias were suppressed by isoflurane anesthesia, Gal OE and WT mice underwent surgical implantation of radiotelemeters for conscious rhythm analysis. Continuous electrocardiograms were recorded before (**Figure 8I**) and after administration of epinephrine 2 mg/kg i.p. (**Figure 8J**). Recordings were extended overnight to capture electrograms during high activity periods. Neither Gal OE nor WT mice had ventricular arrhythmias. Additionally, there was no significant change in the number of PVCs during the study (**Figure 8H**).

**Figure 8.**
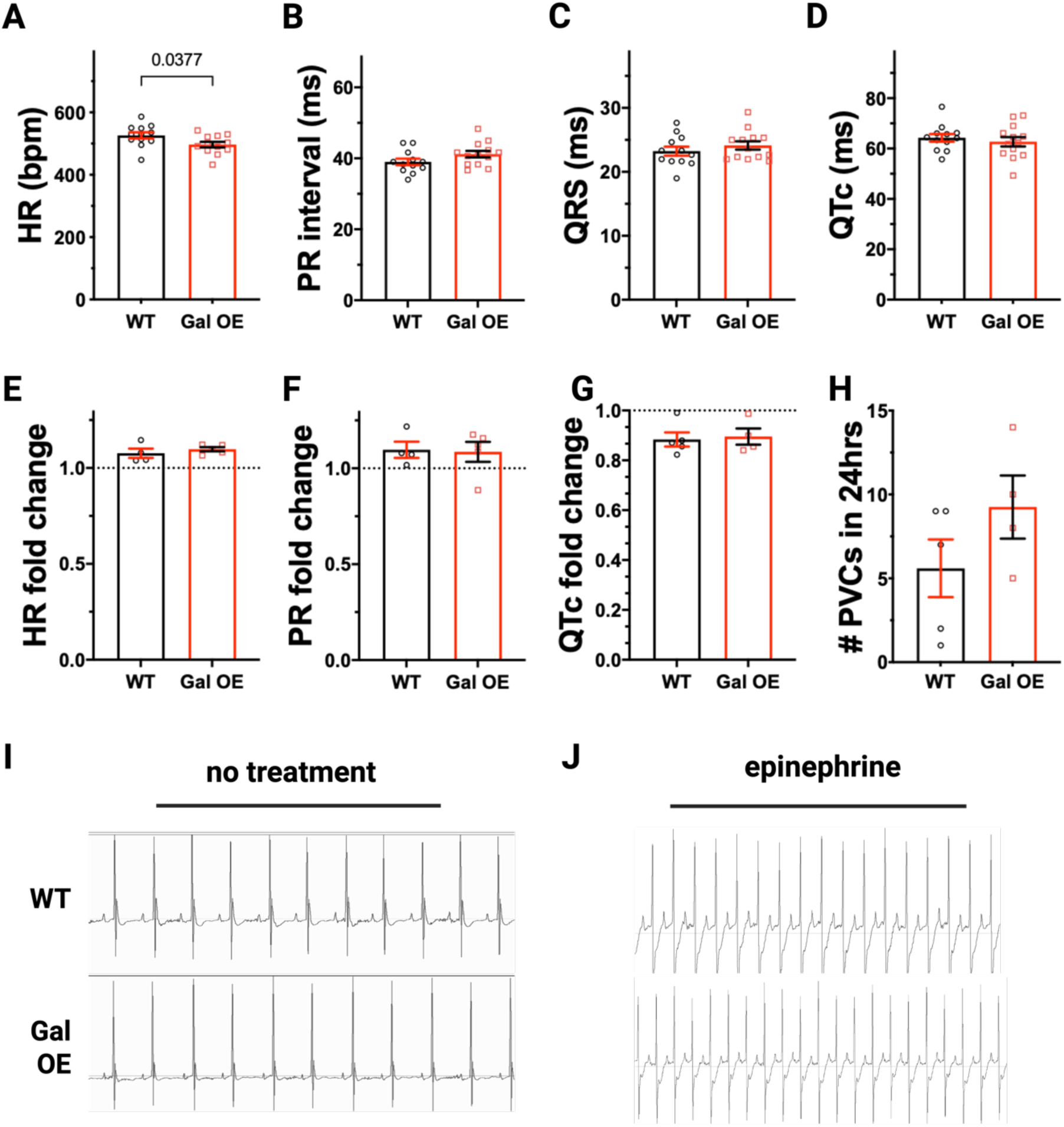
Galectin-1 overexpression does not cause arrhythmia. **A**, **B**, **C**, and **D**. HR, calculated from RR intervals, PR interval, QRS duration, and corrected QT intervals were averaged from 3 random selections on surface electrocardiogram for isoflurane anesthetized WT and Gal OE mice. N=12 WT mice and N=13 Gal OE mice. **E**, **F** and **G**. Changes in HR, PR interval and QTc were the same between groups after treatment with isoproterenol 2 mg/kg i.p. N=4 WT and N=5 Gal OE mice. **H**. PVCs were tallied over a 24-hour period in conscious mice with radiotelemeter insertion after treatment with epinephrine 2 mg/kg i.p. N=5 WT mice and N=5 Gal OE mice. **I** and **J**. Representative electrocardiographic traces of WT and Gal OE mice before and 5 minutes after treatment with epinephrine. Statistical significance was calculated by two-tailed Student’s T-test.

## DISCUSSION

In this study, we applied proximity proteomics to quantitatively define how heart failure (HF) remodels the cardiac dyad—a specialized subcellular domain essential for excitation–contraction coupling and cardiac inotropy, lusitropy, and arrhythmogenesis. Our analysis identified a broad spectrum of proteomic changes in the dyad, providing new mechanistic insights into HF pathobiology. By analyzing myocardium remote from the infarct zone several months post– myocardial infarction, we captured chronic remodeling in a manner that enhances translational relevance to both ischemic and non-ischemic cardiomyopathies.

A key strength of this study lies in the use of two genetically engineered mouse lines expressing APEX2 fused to either the α_1C_ or β_2B_ subunits of Ca_V_1.2—a channel complex that serves as a regulatory hub for cardiomyocyte contractility^45^. Numerous studies have implicated dyadic Ca_V_1.2 dysfunction in HF^8,12,45–47^. By directing proximity labeling through two subunits of Ca_V_1.2, we achieved enhanced spatial resolution and specificity. Although we observed limited differences in the enrichment of canonical dyad proteins such as Ryr2, Jph2, and SERCA2, this likely reflects low tolerance for large-scale changes in their localization and abundance given their essential roles in cardiomyocyte function.

A limitation of the current study is the premature death of several APEX2-α_1C_ and -β_2B_ mice after myocardial infarction, which may have excluded animals with the most profound dyadic remodeling. Additionally, enrichment in proximity proteomics reflects both proximity and protein abundance, introducing interpretative ambiguity. We examined total Psmd4 abundance (all subcellular domains) between Ctrl and HF hearts and found no difference, suggesting there is increased proximity to Ca_V_1.2 rather than increased global expression.

Our findings of dyadic enrichment of microtubule components in HF align with prior work showing expansion of the microtubule network with increase in longitudinally oriented sub-sarcolemmal microtubules in failing myocardium^40^. Given that Ca_V_1.2 surface expression is reduced in HF^48–50^, our enrichment of proteasome components near the channel complex is intriguing and warrants further study. Similarly, our increased enrichment of a protein known to attenuate cardiac dysfunction in ischemic and pressure overload models of HF, Lrrc10, in APEX2-β_2B_ HF hearts also aligns with prior work^38^.

Most notably, we identified Galectin-1 (Gal-1) as a dyad-enriched protein in HF. Gal-1, a β-galactoside-binding lectin with immunomodulatory roles, is highly expressed in muscle tissues and is upregulated in human HF^25,26^. Gal-1 KO mice have reduced systolic function at baseline and increased fibrosis following myocardial infarction, effects attributed to increased inflammation in the heart. Our data, using APEX2 fused to cytoplasmic domains of Ca_V_1.2 subunits, demonstrate endogenous Gal-1 localization within the dyad—a previously unrecognized intracellular site of action.

To investigate the functional implications, we generated mice with inducible, cardiomyocyte-specific overexpression of Gal-1. These mice exhibited increased activation of Ca_V_1.2, RyR2, and SERCA2, consistent with the enhanced inotropy and lusitropy we observed. Phosphorylation of RyR2 (S2808) and Pln at PKA target sites mirror the effects of β-adrenergic stimulation. Our findings suggest an alternative explanation for impaired function in Gal-1 KO mice: reduced Ca^2+^ influx, diminished SR Ca^2+^ load, and impaired systolic function. While global systolic and diastolic function were unchanged by echocardiography in healthy Gal OE and WT mice, we will note that the salutary effects of many proteins are only unmasked under pathologic stress.

Collectively, the impaired function in Gal-1 KO mice, increased cellular contractility in Gal-1 overexpressing mice, upregulation of Gal-1 in human HF, and its dyadic enrichment in failing mouse hearts support the hypothesis that increased dyadic Gal-1 is a compensatory, adaptive response to HF. Isoproterenol markedly increases the Ca^2+^ transient and accelerates relaxation in Gal-1 OE and WT mice alike, suggesting that Ca^2+^ handling proteins in the dyad remain sub-maximally activated and that it may be worth exploring whether or not increasing levels of Gal-1 even higher can improve its inotropic/lusitropic effects. Because of their arrhythmogenic effects, inotropic agents that increase intracellular Ca^2+^ are largely reserved for patients in acute cardiogenic shock or for palliation of symptoms in patients with end-stage disease^1^. Though Gal-1 also increases contractility by augmenting the Ca^2+^ transient, mice with Gal-1 overexpression do not have any greater propensity to adrenergic-mediated arrhythmias.

In conclusion, proximity proteomics revealed Gal-1 as a previously unrecognized dyadic protein with significant effects on Ca^2+^ handling and contractile function in the failing heart. These findings not only refine our understanding of dyadic remodeling in HF but also identify Gal-1 as a potential therapeutic target. The dyad’s capacity to recruit protective proteins such as Gal-1 underscores its role as a dynamic hub in HF pathophysiology.

## ACKNOWLEDGMENTS

Steven Marx, for his gift of transgenic α1C-APEX2 and β2B-APEX2 mice and helpful feedback. Biorender was used for figure preparation.

## SOURCES OF FUNDING

These studies were supported by funding from the National Institutes of Health to JK (K08 HL151969), EW (R01 HL152236), VT (R01 HL170132) and SF (T32 HL120826), the American Heart Association to JSK (Career Development Award 35320208), and the Louis V. Gerstner Scholars Program to JSK. Mice were generated in the Genetically Modified Mouse Model Shared Resource and confocal microscopy images were obtained in the Confocal and Specialized Microscopy Shared Resource of the Herbert Irving Comprehensive Cancer Center at Columbia University, which are supported by the NIH/NCI Cancer Center Support Grant P30CA013696. This work was also conducted with support from the Cancer Prevention and Research Institute of Texas, grant RR220032, to M.K., who is a CPRIT Scholar in Cancer Research.

## DISCLOSURES

No conflicts of interest are noted by the authors.

---

**Supplemental Figure 1.**
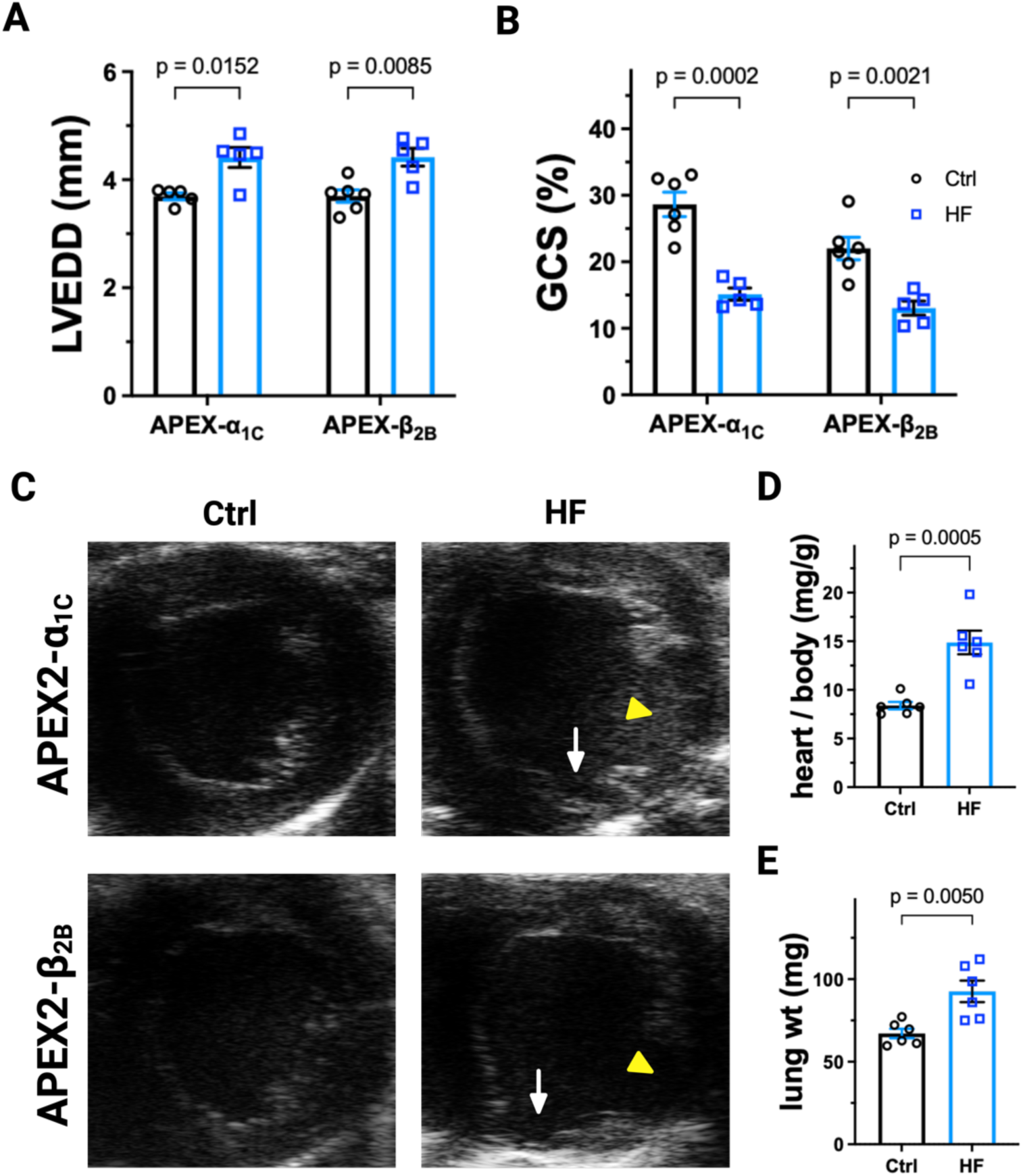
Parameters of heart function in APEX2-α_1C_ and the APEX2-β_2B_ mice after coronary artery ligation. **A**. Left ventricular end-diastolic diameter (LVEDD) was increased in mice in the HF groups. N=6 Ctrl and N=5 HF for both APEX2-α_1C_ and the APEX2-β_2B_ mice. **B**. Global circumferential strain (GCS) was impaired in the HF groups. N=6 Ctrl and N=5 HF for both APEX2-α_1C_ and the APEX2-β_2B_ mice. **C**. Representative B-mode images from parasternal short-axis views at the mid-ventricular level of Ctrl and HF APEX2-α_1C_ and the APEX2-β_2B_ mice White arrows indicate thinned inferior walls and yellow arrowheads indicate LV aneurysm. **D** and **E**. Heart weight to body weight and dry lung weights were measured in APEX2-α_1C_ mice 8 weeks after coronary artery ligation. N=6 Ctrl and N=HF mice. Indicated P-values were calculated by two-tailed Student’s T test.

**Supplemental Table 1.**
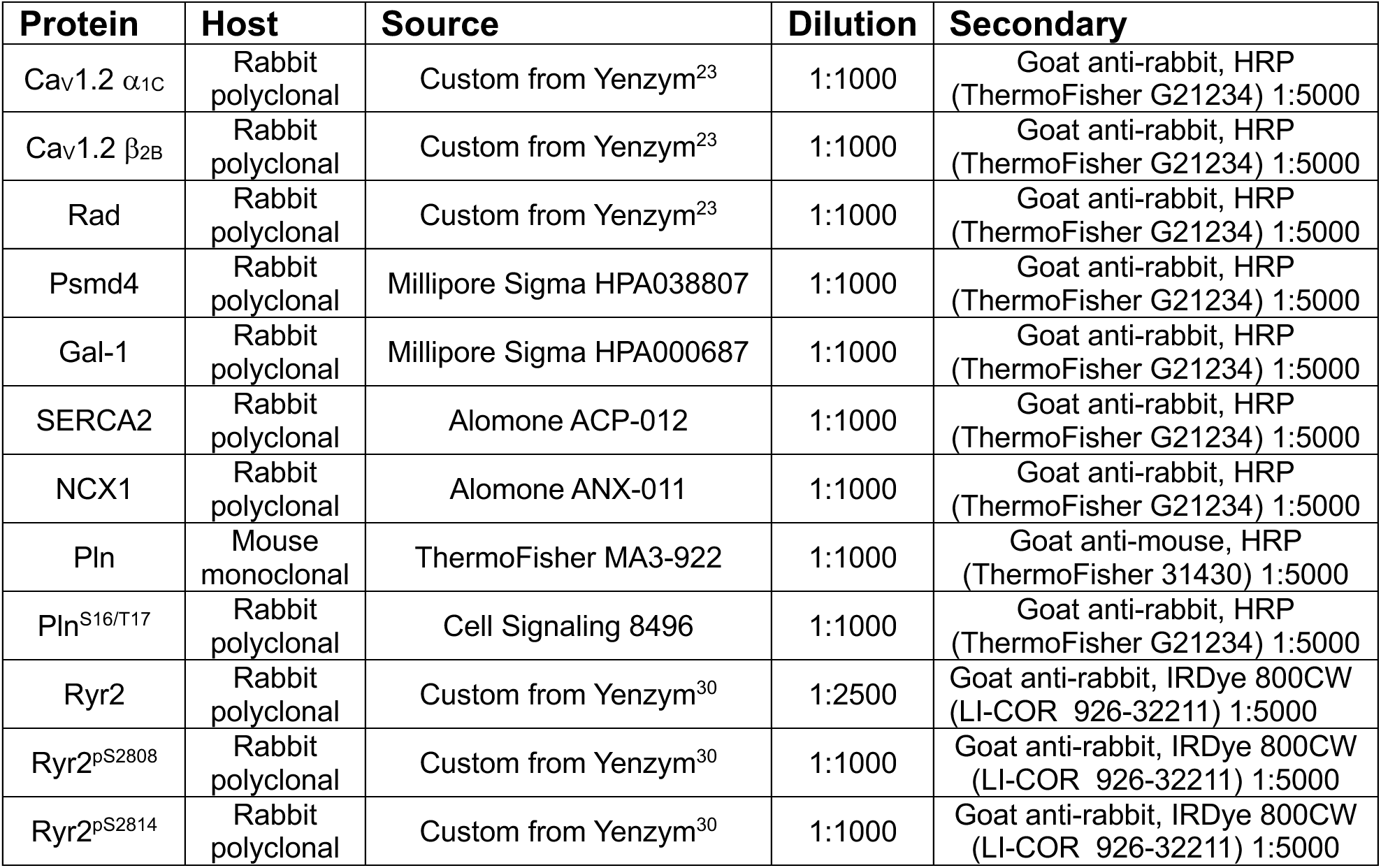
List of antibodies used.

## Notes

### Competing Interest Statement

The authors have declared no competing interest.

